# Poor sleep impairs immune responses and influenza vaccine protection

**DOI:** 10.1101/2025.05.07.652783

**Authors:** Minhui Guan, Weihong Gu, Khalyfa Abdelnaby, Pradeep Balamalaliyage, Wikanda Tunterak, Moran Li, Yuhan Wen, John Driver, David Gozal, Xiu-Feng Wan

## Abstract

Disrrupted sleep, a common occurrence among shift workers, older adults, and individuals with sleep disorders, is increasingly recognized as a potential factor interfering with vaccine efficacy. Using a mouse model, we show that two weeks of chronic sleep fragmentation (CSF) before and during influenza vaccination significantly impair humoral immunity and reduce protection against lethal viral challenge. CSF-exposed mice exhibited lower neutralizing antibody titers, diminished IgG subclass responses, and reduced survival after viral challenge, despite preserved antibody binding avidity. Single-cell RNA sequencing and immune receptor profiling revealed altered B cell maturation, abnormal germinal center activation, and plasma cell stress, characterized by activation of unfolded protein response and oxidative stress pathways. CSF also reprogrammed B-cell signaling and disrupted coordination with T-cells. Together, this study showed that CSF compromises vaccine-induced immune responses by affecting multistage of B-cell differentiation, underscoring the importance of considering sleep health in vaccination strategies for vulnerable populations.

**One Sentence Summary:** Poor sleep weakens both the magnitude and quality of immune responses, compromising the protective efficacy of influenza vaccination.

## Introduction

Since 2010, annual influenza vaccination has been recommended for persons aged six months and older (*1*) and vaccination remains the primary strategy for limiting the global influenza burden (*2*). However, influenza vaccine effectiveness varies dramatically among individuals due to factors such as pre-existing immunity (*3–12*), age (*13–15*), sex (*16*), host genetic polymorphisms (*17, 18*), pre-existing health conditions (e.g. obesity and diabetes) (*19*), socio-economic status, and other variables (*20*).

Chronic sleep fragmentation (CSF), which affects approximately one-third of the global population (*21*), is a major form of sleep disruption that is characteristically present among shift workers, healthcare professionals, parents of young children, frequent travelers, and elderly adults who often experience sleep discontinuity. Additionally, individuals with sleep disorders such as sleep apnea, restless legs syndrome, or insomnia, those suffering from chronic pain or medical conditions (e.g. cancer, chronic back pain, fibromyalgia, gastroesophageal reflux disease, cancer, obesity, diabetes, cardiovascular diseases, recurrent infections, and depression) (*22*), and those exposed to environmental noise, are also significantly affected. Patients with CSF are at increased risk of developing excessive daytime sleepiness (*23*) and various end-organ morbidities, particularly impacting cardiovascular (*24*), metabolic (*25*), cognitive, and behavioral functioning (*26*). The prevalence and severity of CSF increase with age (*27*). Thus, CSF is expected to become an increasing problem in aging populations around the world (*28*).

Sleep deprivation and poor sleep quality have been linked to alterations in both innate and adaptive immune responses, chronic inflammation, and increased infection risk (*29*). Prolonged wakefulness can elevate leukocyte counts in the blood (e.g., white blood cells, lymphocytes, monocytes, B-cells, and neutrophils) and increase pro-inflammatory cytokines such as IL-1, IL-6, and TNF (*30*). Additionally, adequate sleep is essential for maintaining the balance between Th1 and Th2 cytokines (*31*), and is associated with a reduced infection risk, improved disease outcomes, and stronger vaccine responses, including those for influenza, hepatitis A and B vaccines (*32–37*). However, most studies on vaccine efficacy in humans associated with sleep deprivation, including those focused on influenza vaccines, are observational, and the underlying molecular mechanisms are not fully understood.

In this study, we use a well-established murine model to demonstrate that two weeks of sleep disruption before and during influenza vaccination impairs the magnitude and quality of antibodies induced by an influenza vaccine and decreases protection from a subsequent live virus challenge. Through comprehensive analyses of the antibody responses and single-cell transcriptomics data, we uncover the molecular signatures associated with CSF-induced immune impairments.

## Results

### Chronic sleep fragmentation impairs influenza vaccine protection

To evaluate whether CSF affects the protective efficacy of influenza vaccination, we assessed immune responses, morbidity, and mortality in sleep-disrupted and normal-sleep control C57BL/6J mice following vaccination and challenge with live influenza virus. To ensure that the immunological effects of CSF could be sensitively and reproducibly detected, we implemented a carefully optimized vaccine dosing and viral challenge strategy. Our goal was to establish immunization conditions that elicited robust, yet submaximal protection, sufficient to prevent complete lethality while rendering mice susceptible to modulation by physiological stressors such as CSF. This approach allowed us to identify impairments in vaccine-induced immunity and increased mortality specifically attributable to CSF exposure.

In this study, we tested immune responses to vaccine doses of 0.1, 0.2, and 0.3 μg HA and found that all doses generated detectable vaccine strain–specific titers by both hemagglutination inhibition and microneutralization assays (**Supplementary Fig. 1a**). Following challenge with 500 LD₅₀, well-rested mice vaccinated with 0.2, 0.3, or 1 μg HA were fully protected against lethal challenge doses of up to 5,000 LD₅₀ (**Supplementary Fig. 1b**). In two independent experiments with mice vaccinated using 0.1 μg HA, one experiment showed partial protection with an 80% survival rate against a 500 LD₅₀ challenge, while the other achieved full protection, suggesting that the 0.1 μg HA–500 LD₅₀ combination represents the threshold for complete protection in well-rested mice. These results are consistent with a previous study demonstrating that prime–boost immunization with as little as 0.01–0.02 μg of HA protein could elicit strong H1-specific IgG responses (∼200 μg/mL) and confer full protection against homologous H1N1 challenge, while doses as low as 0.005 μg provided partial protection (*38*). Such reports highlight the remarkable sensitivity of mouse models to low-doses of HA immunization, which can provide protection at HA concentrations that are far lower than the typical 1–15 μg range used in other studies. By leveraging this low-dose, high-resolution model, we aimed to ensure that the system was sufficiently sensitive to detect the subtle but biologically meaningful effects of CSF on humoral immunity and vaccine efficacy.

Based on these results from our dose optimization experiments, we selected a 0.2 μg HA vaccine dose and a 500 LD₅₀ challenge dose for our main experiment. A total of 105 six- to eight-week-old male mice were separated into four experimental groups: Groups 1 and 2 were subjected to either CSF (gSF) (n = 30) or normal-sleep control (gSC) (n = 30), respectively. Both groups were then vaccinated with whole inactivated A/California/07/09 (H1N1) (CA/07) vaccine and challenged with antigenically identical, mouse-adapted A/California/04/09 (H1N1) (CA/04-MA). Group 3 (Mock; n = 15) consisted of normal-sleep mice sham-vaccinated with PBS and challenged with CA/04-MA. Group 4 (Negative; n = 30) consisted of normal-sleep mice sham-vaccinated and sham-challenged with PBS (**Fig. 1a**). CSF exposure consisted of mechanical sleep interruptions every two minutes during the 12-hour lights-on phase each day corresponding to the usual rest period in rodents (**Fig. 1b**). The CSF exposure began two weeks before vaccination and continued through vaccination and viral challenge phases.

**Fig. 1.**
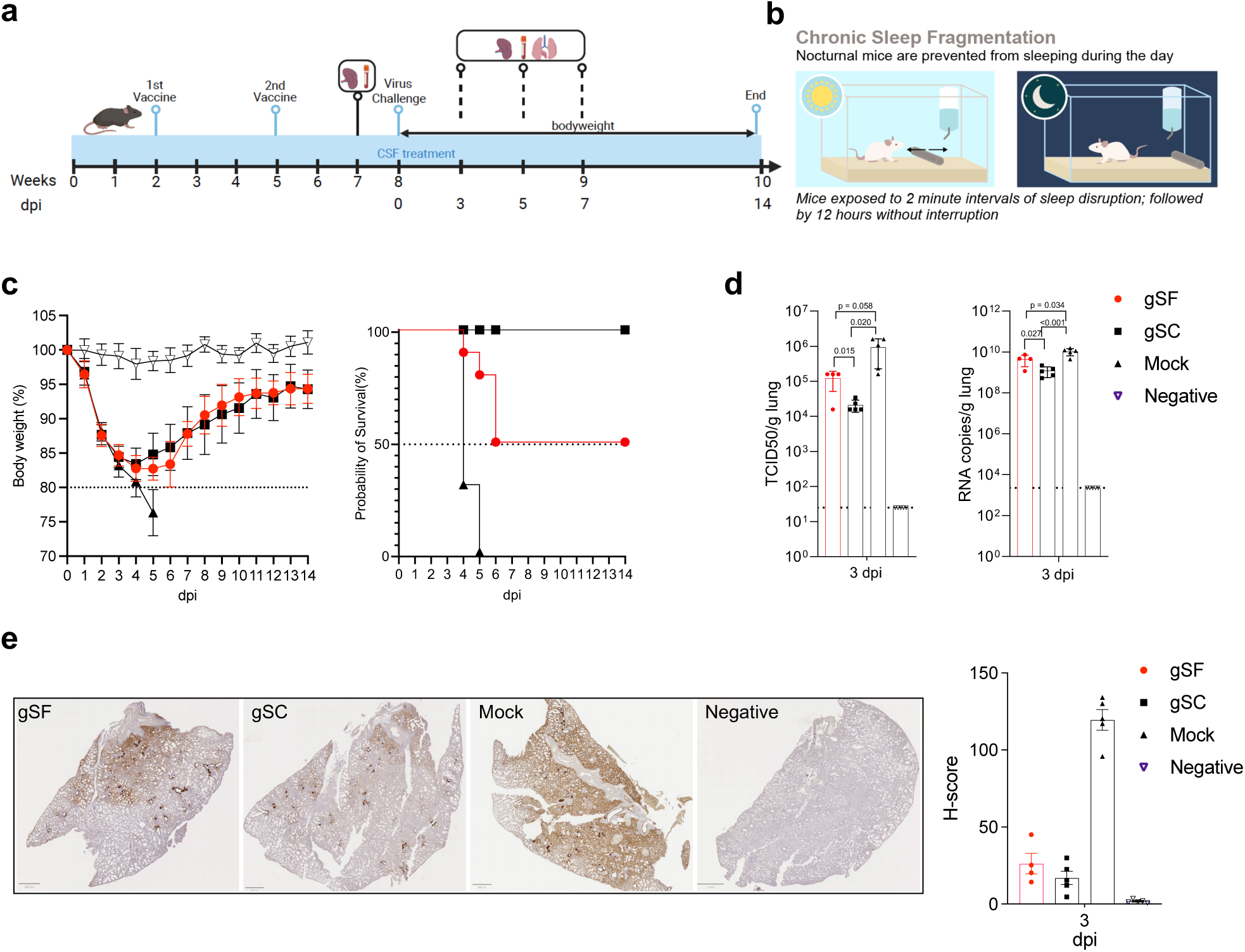
Chronic sleep fragmentation (CSF) impairs influenza vaccine protection. **a)** Experimental design. A total of 105 six- to eight-week-old male C57BL/6J mice were assigned to four groups: gSF (SF+ vaccine+virus challenge; n=30), gSC (sleep control+vaccine+virus challenge; n=30), Mock (sleep control+PBS+virus challenge; n=15), and Negative (sleep control+PBS+PBS; n=15). Each mouse received a vaccination dose of 0.2 µg HA of A/California/07/09 (H1N1) (CA/07) and a challenge dose of 500 LD_50_ of mouse-adapted A/California/04/09 (H1N1) (CA/04-MA) via intranasal inoculation when required. Both doses for vaccination and virus challenges were optimized (**see Supplementary Fig. 1**). Of note, CA/07 and CA/04-MA are antigenically identical. Five mice per group were sacrificed before challenge and at 3, 5, and 7 days post infection (dpi) for immunological and pathological assessments. Remaining mice were monitored through 14 dpi. **b)** Design of CSF treatment. CSF treatment consisted of sleep interruptions every 2 minutes for 12 hours/day, followed by 12 hours of undisturbed rest, beginning 2 weeks before vaccination and continuing through the study. **c)** Dynamics of body weight changes and mortality after lethal challenges: Body weight of live animals and survival rates of the mice were monitored. The experiments were repeated four times (**See Supplementary Fig. 1**). **d)** Viral loads in the lung at 3 dpi after lethal challenges. Infectious virus titers were determined by TCID₅₀ assay and expressed as log₁₀ TCID₅₀/gram tissue. Viral RNA levels were quantified by M gene–specific quantitative RT-PCR and expressed log₁₀ copies/gram tissue. Dashed lines indicate the limits of detection: 10^1.69^ TCID₅₀/gram and 10^3.65^ viral RNA copies/gram. **e)** Distribution of influenza antigens and NP staining scoring. Quantification of influenza NP-positive cells using QuPath 0.4 (*71*) across the entire slide. H-scores were measured from five mice lung per group, except for the gSF group, which included only four mice due to the loss of one animal. All samples were de-identified before analysis to ensure blinding.

CSF-exposed mice exhibited greater than 80% weight loss and a 50% survival rate after lethal challenge, whereas all vaccinated control (gSC) mice survived the challenge (**Fig. 1c**). As anticipated, all mice in the sham-vaccinated group succumbed to infection, while all mice in the negative control group remained healthy. Lung viral loads at 3 days post-infection (dpi) were significantly higher in CSF-exposed vaccinated mice compared to vaccinated normal-sleep controls, although both groups had lower viral loads than the unvaccinated mock group (**Fig. 1d**). Immunohistochemistry revealed higher concentrations of viral nucleoprotein (NP) in the lungs of CSF-treated mice compared to normal-sleep controls at 3 dpi (**Fig. 1e**). Viral titers in surviving mice gradually declined and became undetectable by 14 dpi.

To validate these findings, we conducted three additional independent experiments using different vaccine and challenge doses. Among CSF-exposed mice, vaccination with 0.1 μg HA followed by a 500 LD₅₀ challenge resulted in a 40% survival rate in two experiments, while vaccination with 0.3 μg HA followed by a 5,000 LD₅₀ challenge resulted in a 50% survival rate (**Supplementary Fig. 1c**). In all replicate experiments, well-rested vaccinated mice achieved complete protection across the tested doses.

Taken together, these results demonstrate that CSF consistently and significantly impairs the protective efficacy of influenza vaccination. Across multiple independent experiments, CSF exposure was associated with increased mortality, elevated lung viral loads, and greater viral antigen burden despite otherwise protective vaccination. These findings demonstrate robust and reproducible impairment of vaccine-induced immunity under CSF.

### CSF reduces the magnitude of influenza vaccine-specific antibody responses

We hypothesized that CSF would impair both the magnitude and quality of vaccine-induced humoral responses. Thus, we assessed serological titers in the sera collected at 35 days post-vaccination (dpv) from mice subjected to CSF (gSF) or normal-sleep control (gSC) conditions. gSF mice had significantly lower hemagglutination inhibition (HAI) titers compared to gSC mice (p < 0.05; **Fig. 2a**). Neutralizing antibody titers, measured by microneutralization (MN) assays, were also reduced in CSF-exposed mice, although the difference was not statistically significant (p = 0.0877). In contrast, IgG avidity, assessed using both recombinant HA proteins and the whole vaccine virus CA/07, was similar in gSF and gSC groups (**Fig. 2b**), suggesting that CSF primarily affects antibody quantity rather than binding strength.

**Fig. 2.**
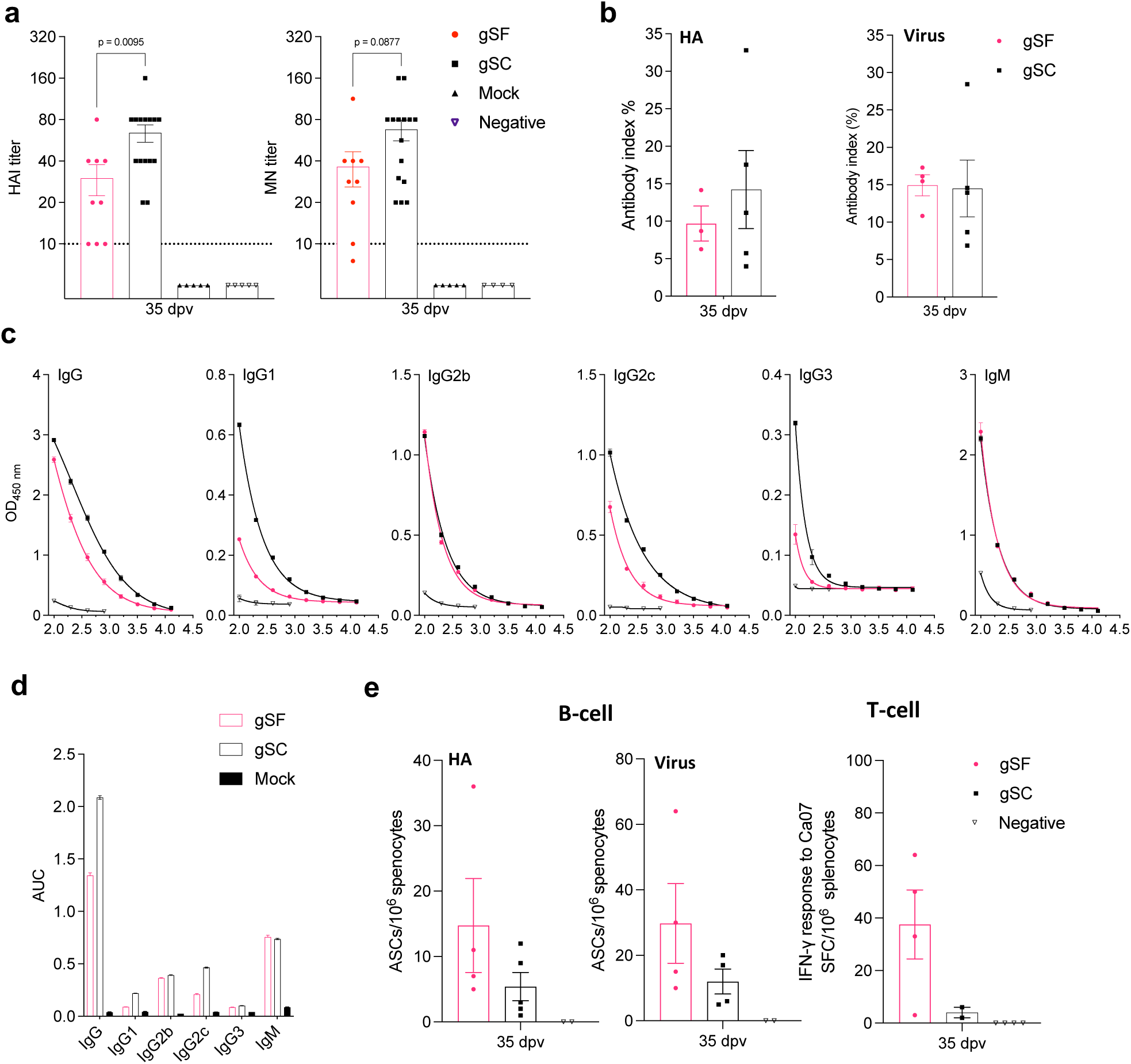
Chronic sleep fragmentation reduces influenza vaccine specific humoral responses. **a)** Serological titers post-vaccination. Serological titers in the mice blood at 35 days post-vaccination (dpv) but before challenge, were determined by hemagglutination inhibition (HAI) and microneutralization assays. **b)** Avidities of vaccine-specific antibodies. The avidity index was calculated based on the area under the entire antibody titration curve using GraphPad Prism (version 10.1.2). The antibody index equals to an AUC ratio of treated and untreated reactions. All samples were assessed in duplicate, and mean titers were calculated (*74*). **c)** quantities of total IgG and subclass IgGs. **d**) Aera under curves (AUC) for total IgG and subclass IgGs. **e)** ELISpot analysis of HA- and vaccine-specific antibody-secreting B-cells and virus-specific IFN-γ–producing T-cells. Splenocytes were collected from CSF-treated (gSF) and normal-sleep control (gSC) mice one week prior to viral challenge. The frequency of antigenic-specific IgG-secreting B-cells was assessed by ELISpot following in vitro stimulation with recombinant HA protein of CA/07 and the purified vaccine virus. The frequency of virus-specific IFN-γ–producing T-cells was measured by ELISpot after stimulation with whole CA/07 virus. Data are shown as the number of spot-forming cells per million splenocytes.

To further characterize humoral responses, we quantified total vaccine-specific IgG and analyzed IgG subclasses in the sera collected at 35 dpv. Total IgG, along with IgG1, IgG2c, and IgG3 subclasses, were significantly reduced in CSF-treated mice (**Fig. 2c and 2d**). In contrast, IgG2b and IgM levels were similar between groups. Notably, in humans, IgG subclasses IgG1 and IgG2 are typically elevated after influenza vaccinations (*39, 40*). In the mouse model, simultaneous induction of IgG1 and IgG2c reflects a balanced Th2- and Th1- immune response that is often associated with superior protective immunity (*41*). Of note, in C57BL/6J mice, IgG2c serves as the functional equivalent of IgG2a, which is not expressed in this strain (*42*).

Interestingly, ELISpot assays showed comparable or slightly higher frequencies of influenza-specific antibody-secreting B-cells and IFN-γ–producing T-cells in the spleens of CSF-treated mice compared to well-rested controls at 35 dpv (**Fig. 2e**). These findings suggest that the reduced influenza-specific antibody concentrations in CSF-treated mice were not due to lower numbers of antigen-specific lymphocytes. Instead, CSF likely does not impair the initial priming of these cells but rather disrupts their functional differentiation or effector output, resulting in diminished humoral immunity, despite preserved or even elevated cellular frequencies.

Together, these findings demonstrate that CSF disrupts vaccine-induced humoral immunity by reducing antibody titers and altering IgG class switching, but without reducing the concentration of virus-specific T and B lymphocytes.

### CSF reprograms immune transcriptional states associated with humoral impairment

To define the cellular and molecular basis underlying impaired vaccine responses in CSF-treated mice, we performed single-cell RNA sequencing (scRNA-seq) and paired T-/B-cell receptor (TCR/BCR) sequencing on splenocytes collected at 35 dpv, one week prior to viral challenge. Splenocytes (∼10⁴ cells/mouse) from three gSF and three gSC mice were processed using the Chromium Next GEM Single Cell 5’ Kit v2 (10x Genomics) and sequenced on an Illumina NovaSeq platform.

Unsupervised clustering identified 25 transcriptionally distinct cell populations, annotated based on canonical lineage markers (**Fig. 3a and Supplementary Fig. 2a**). Among these, 24 clusters corresponded to immune cells, including eleven B-cell clusters, six T-cell clusters, and seven innate immune cell clusters, including NK cells, macrophages, conventional dendritic cells (cDC1), monocyte-derived dendritic cells (MoDCs), plasmacytoid dendritic cells (pDCs), neutrophils, and mast cells. The integration of gSF and gSC samples revealed that both groups had comparable cell cluster frequencies (**Fig. 3b and Supplementary Fig. 2b**). To assess the transcriptional impact of CSF, we quantified differentially expressed genes (DEGs) within each immune subset. DEG counts varied markedly across clusters (**Fig. 3c**), with pronounced transcriptional differences observed in neutrophils (Cluster 23), mast cells (Cluster 24), MoDCs (Cluster 21), macrophages (Cluster 19), and double-negative (DN) T-cells (Cluster 16). Among B-cell subsets, immature B-cells (Cluster 1), germinal center (GC) B-cells (Cluster 10), and plasma cells (Cluster 11) exhibited the greatest number of CSF-induced transcriptional changes.

**Fig. 3.**
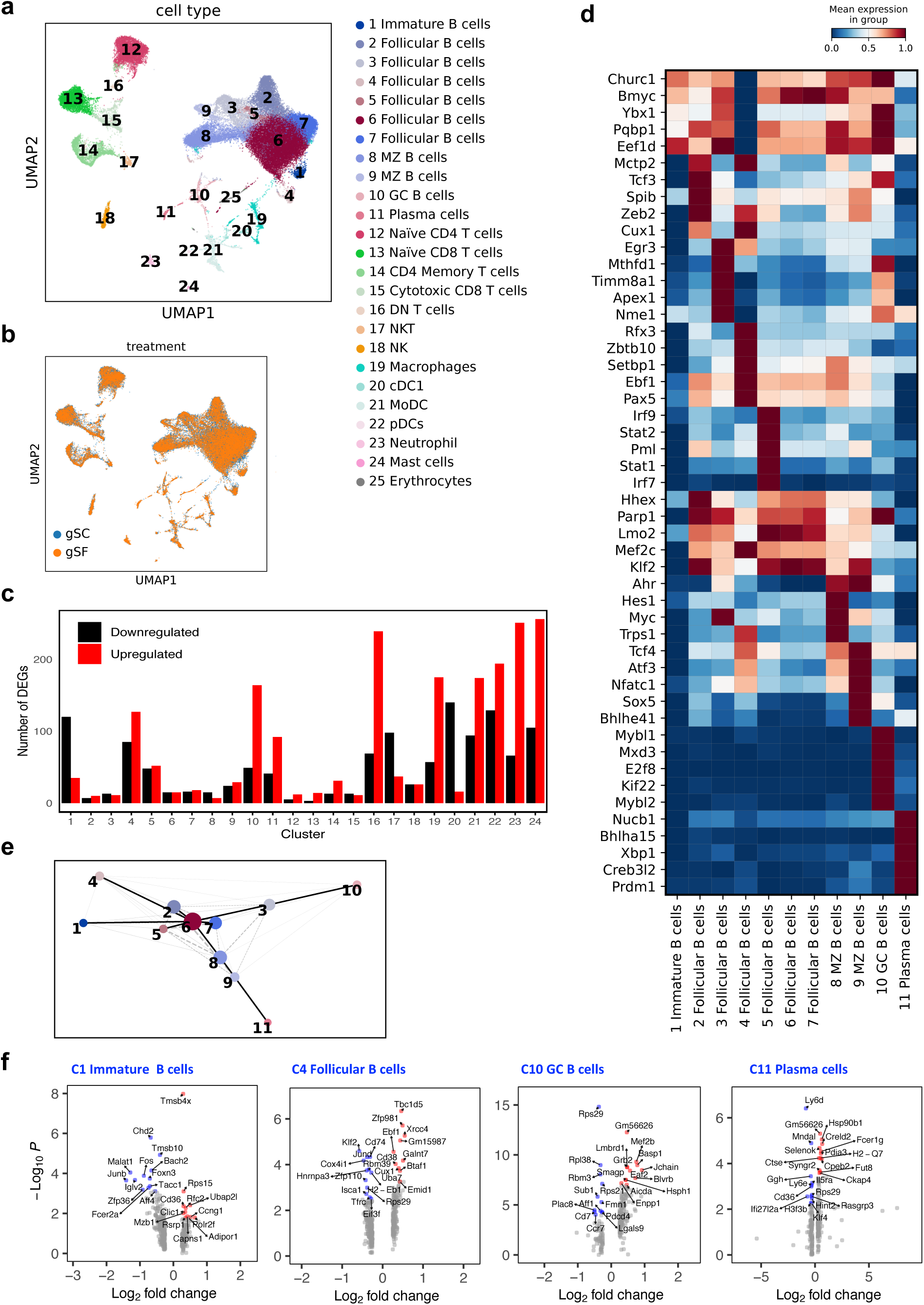
Single-cell transcriptomic profiling of spleen cells from mice exposed to Chronic sleep fragmentation. **a)** Uniform Manifold Approximation and Projection (UMAP) projection of spleen cells from CSF- and normal sleep-exposed mice. UMAP was used to visualize single-cell RNA sequencing data. Distinct cell clusters are color-coded to represent different immune cell types or states. **b)** UMAP plot of integrated gSF and gSC groups. **c)** Bar plot showing the number of differentially expressed genes (DEGs) with p-value < 0.05 per cell type or cluster for immune cells. **c)** Heatmap of the top five transcription factors for each B-cell cluster. **d)** Partition-based graph abstraction (PAGA) analysis illustrating the transcriptional connectivity and lineage relationships among B-cell subsets. **e)** Volcano plots showing differentially expressed genes between gSF and gSC. Red and blue dots represent the top upregulated and downregulated genes in gSF compared to gSC, respectively.

Analysis of transcription factor activity across B-cell subsets revealed that GC B-cells and plasma cells form transcriptionally distinct states relative to other B-cell populations (**Fig. 3d**). A pseudotime trajectory analysis of B-cell subsets support a branching differentiation model in which immature (Cluster 1) progress through follicular B-cells (Clusters 2–7) before diverging into marginal zone (Clusters 8–9), GC (Cluster 10), or plasma cell (Cluster 11) fates (**Fig. 3e**).

Given the observed reduction in antibody responses, we examined four representative B-cell subsets in greater detail (**Fig. 3f**). Immature B-cells had a substantial number of up and down regulated genes, such as *Tmsb4x* and *Cd36* that were upregulated, and the key transcriptional regulators including *Foxn3*, *Fos*, *Junb*, and *Bach2*, which were downregulated. A population of follicular B-cells (FoB), which we designated as activated FoB cells (**Supplementary Fig. 3a**), exhibited moderate shifts in gene expression, including increased *Ebf1*, a transcription factor involved in B-cell lineage programming and identity maintenance (*43*), and decreased expression of *Cd74*, a key chaperone for MHC class II antigen presentation (*44*), *Cd38*, a marker of B-cell activation and germinal center transition (*45*), and *Rbm39*, an RNA-binding protein involved in transcriptional regulation and immune cell function (*46*). GC B-cells showed a robust transcriptional response, with marked upregulation of *Aicda*, *Jchain*, and *Blvrb*, which are genes that are essential for antibody class switching and plasma cell transition (*47*), and concurrent downregulation of *Rps29*, *Rpl38*, and *Rbm3*. These changes suggest that CSF impacts protein synthesis and stress regulation. Plasma cells from CSF-exposed mice upregulated genes involved in ER protein folding and stress adaptation, *Pdia3* (oxidative folding) (*48*), *Ckap4* (ER homeostasis) (*49*), and *Syngr2* (vesicle trafficking) (*50*). Conversely, they downregulated *Ifi27l2a* (interferon response) (*51*), *Rps29* (hematopoietic dysfunction and ribosomal stress responses) (*52*), and *Cd36* (lipid metabolism and immune regulation) (*53*), supporting a model of ER stress adaptation and reduced effector function.

Together, these findings demonstrate that CSF alters B-cell developmental trajectories and reprograms key transcriptional programs required for effective humoral immunity.

### CSF alters B-cell signaling programs and disrupts coordination with T-cells

To investigate functional consequences of the transcriptional reprogramming observed in B-cells, we performed Ingenuity Pathway Analysis (IPA) on DEGs between gSF and gSC B-cell clusters (**Fig. 4a**). In immature B-cells, CSF exposure downregulated multiple pathways associated with early activation, oxidative stress regulation, and cellular metabolism, including BCR signaling, NRF2-mediated stress response, sirtuin signaling, and retinoic acid receptor (RAR) pathways. Follicular and marginal zone B-cells exhibited relatively few pathway-level changes. In contrast, plasma cells showed broad activation of pathways involved in antigen presentation (MHC class II), BCR signaling, the unfolded protein response (UPR), and cellular stress adaptation, including protein ubiquitination, HMGB1 signaling, and chaperone-mediated autophagy. Germinal center (GC) B-cells and plasma cells also demonstrated enrichment of immunoglobulin class switching programs, including NFAT signaling and interferon gamma responses, indicating enhanced engagement in effector differentiation.

**Fig. 4.**
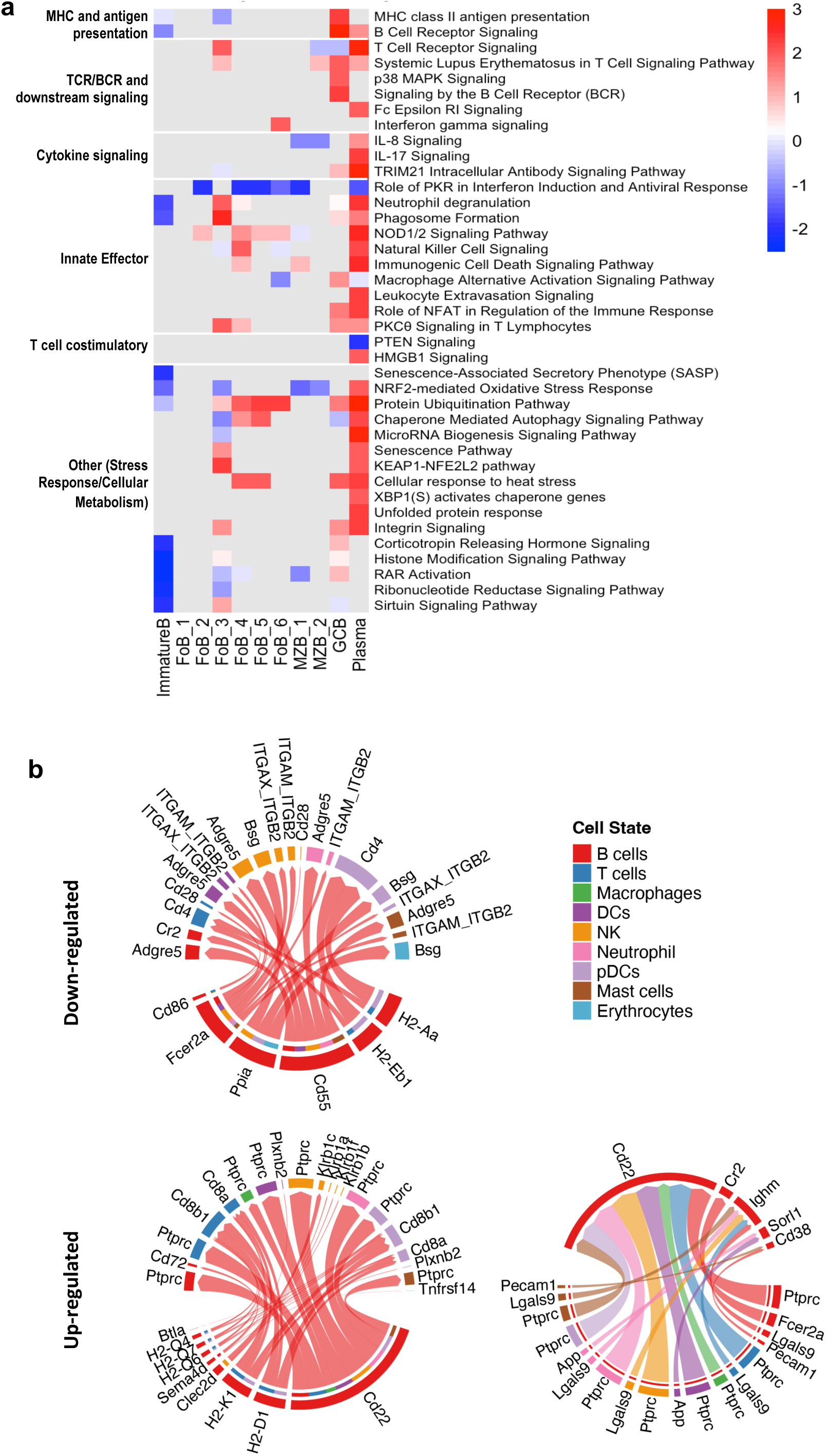
Pathway analysis and disrupted B–T cell coordination under chronic sleep fragmentation. **a)** Ingenuity Pathway Analysis (IPA) of DEGs in B-cell subsets. Core Analysis function was used to identify canonical pathways significantly associated with the DEGs, applying a p-value threshold of 0.05. Pathways were considered significant with –log(p-value) > 1.3 (p < 0.05) and an activation Z-score > 2 or < –2. Only pathways with Z-scores ≥ 2 or ≤ –2 in at least one B-cell type were included (performed using R). Red indicates pathways upregulated in CSF mice. Pathways were grouped into six categories: 1) MHC and antigen presentation, 2) TCR/BCR and downstream signaling, 3) Cytokine signaling, 4) Innate effector functions, 5) T-cell costimulatory signals, 6) Other such as stress response and cellular metabolism. **b)** CSF alters B–T cell communication by downregulating antigen presentation and adhesion pathways while enhancing costimulatory signals. Chord diagrams depict ligand–receptor interactions between B-cells and T-cells inferred from single-cell RNA-seq using CellChat, based on differentially expressed genes (DEGs) in CSF-exposed (gSF) mice compared to normal-sleep control (gSC) mice. Top panel: Downregulated ligand–receptor interactions in CSF mice. Key pathways diminished under CSF include antigen presentation (e.g., H2-Aa–Cd4, H2-Eb1–Cd4) and adhesion (e.g., ICAM1–LFA1, Cr2–Cd21, Fcgr2b– Cd32), reflecting impaired immunological synapse formation and reduced coordination between B- and T-cells. Bottom left panel: Upregulated ligand–receptor interactions in CSF mice. These include enhanced costimulatory signaling between B- and T-cells (e.g., Cd86–Cd28, Cd80–Cd28, ICAM1–LFA1), suggesting increased demand for T-cell help but in the absence of sufficient antigen presentation. Bottom right panel: Summary comparison of the altered B–T communication landscape in CSF mice, highlighting an imbalance where costimulatory signals are elevated but antigen presentation and adhesion pathways are suppressed, pointing to a decoupling of activation and specificity under chronic sleep disruption.

Innate immune subsets were also affected. Macrophages, dendritic cells, and neutrophils exhibited coordinated upregulation of genes involved in antigen processing, oxidative stress, phagosome formation, and ER stress signaling (**Supplementary Fig. 2c**). In contrast, most conventional T-cell subsets were transcriptionally stable. One exception was the DN T-cell cluster, which displayed upregulation of pathways involved in T-cell receptor signaling, CTLA-4 signaling, ferroptosis, and autophagy (**Supplementary Fig. 2c**).

To evaluate whether CSF disrupted immune coordination at the intercellular level, we applied CellChat (v2.1.1) (*54*) to infer ligand–receptor interactions across immune compartments. In gSF mice, B cells upregulated several key ligands, including *Cd22* and MHC class I molecules, which are predicted to engage *Ptprc* (encoding CD45), and the CD8 molecules *Cd8a* and *Cd8b1, respectively,* on T cells and other immune subsets (**Fig. 4b**). This upregulation suggests enhanced demand for T cell help and heightened immune activation (*55*). However, gSF B cells also showed reduced ligand expression of costimulatory ligand Cd86, *H2-Aa*, a key MHC class II molecule required for antigen presentation to CD4⁺ T cells, and *Fcer2a* (CD23, a low-affinity receptor for IgE), which is involved in regulating IgE levels as well as B cell growth and differentiation (*56*). Notably, significantly downregulated receptor signaling networks were not observed in the gSF condition B cells.

Collectively, these findings reveal that CSF exposure reconfigures signaling pathways within B-cells and alters their coordination with T-cells. The combined upregulation of CD8 T cell-B cells signaling and downregulation of CD4 T cell-B cells interactions suggests a decoupling of immune activation and specificity, which may compromise adaptive immune precision and efficiency under sleep-disrupted conditions.

### CSF alters TCR and BCR diversity and repertoire architecture

To evaluate whether CSF alters immune receptor selection of T- and B-cells, we performed paired TCR and BCR sequencing on splenocytes from gSF and gSC mice. A UMAP projection of all cells with annotated immune receptor identities revealed distinct clustering of BCR+ and TCR+ cells with minimal overlap, confirming accurate lineage assignment (**Fig. 5a**).

**Fig. 5.**
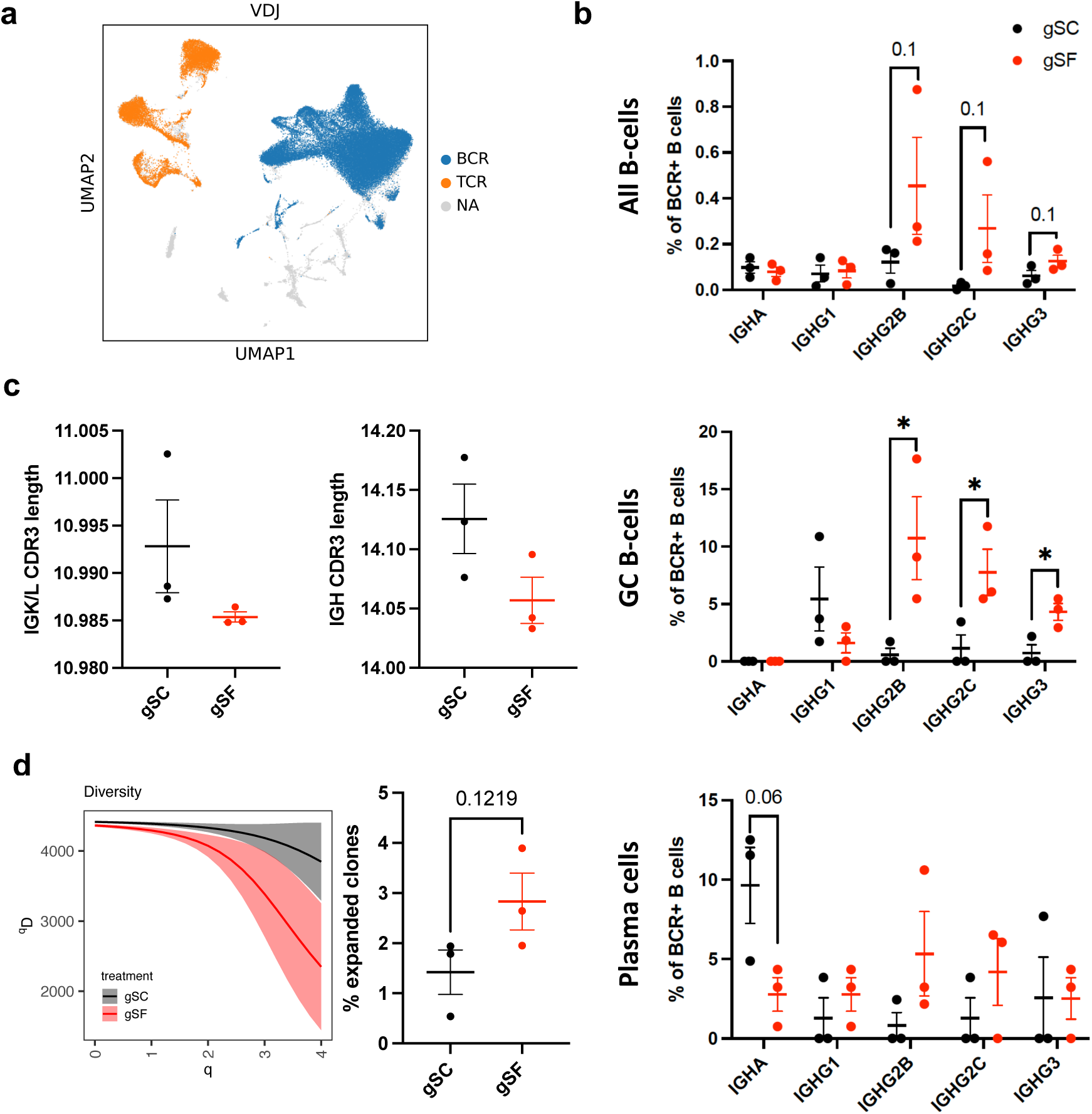
B-cell and T-cell repertoire diversity under chronic sleep fragmentation. **a)** UMAP projection of splenocytes expressing productive TCR or BCR chains, highlighting distinct clustering of BCR⁺ and TCR⁺ cells in CSF-exposed (gSF, red) and control (gSC, black) mice. **b)** Immunoglobulin isotype distribution among BCR⁺ B-cells within total B-cells (top), germinal center (GC) B-cells (middle), and plasma cells (bottom). **c)** Mean length distributions of complementarity-determining region 3 (CDR3) of immunoglobulin. Each dot represents an individual mouse. **d)** Clonal diversity of TCR repertoires (left) and clonal expansion of TCR repertoires (right). Hill numbers quantify diversity: q = 0 reflects clonal richness (equal weighting), q = 1 corresponds to Shannon entropy (moderate sensitivity to evenness), and q = 2–4 emphasize the dominance of expanded clones. qD represents the effective number of clones for a given q. Each dot represents an individual mouse; bars denote mean ± SD. Statistical comparisons were made using unpaired two-tailed t-tests; *p* < 0.05 was considered significant.

To investigate the impact of CSF on B-cell diversification, we analyzed immunoglobulin isotype usage across transcriptionally defined B-cell clusters. Class-switch recombination (CSR) was enriched in GC B-cells, which showed increased expression of IgG2b, IgG2c, and IgG3 (p < 0.05), and in plasma cells, which expressed higher levels of IgA (p = 0.06) (**Fig. 5b**). Other clusters remained dominated by IgM, consistent with naïve or early-stage B-cells (**Supplementary Fig. 3**). We also assessed complementarity-determining region 3 (CDR3) lengths in immunoglobulin heavy (IGH) and light (IGK/IGL) chains. gSF mice exhibited slightly shorter CDR3s compared to gSC mice (**Fig. 5c**). Because CDR3 length influences antigen-binding specificity and is shaped by germinal center selection and somatic hypermutation (*57*), this subtle contraction may reflect impaired repertoire breadth or constrained affinity maturation. While longer CDR3s have been associated with polyreactivity and lower specificity (*58*), shorter CDR3s may limit the diversity of vaccine-induced antibody responses.

In parallel, we compared TCR clonal expansion between gSF and gSC mice. Although the difference did not reach statistical significance, gSC mice exhibited a lower clonal diversity, and consistently, a trend toward greater expansion of dominant T cell clones was observed in gSF mice (**Fig. 5d**). This pattern may reflect subtle alterations in antigen-driven proliferation or persistence of effector T-cell clones under CSF conditions.

Together, these findings suggest that CSF alters the architecture of both TCR and BCR repertoires. Although major developmental processes such as class switching and clonal expansion were preserved, CSF was associated with modest reductions in BCR CDR3 length, and a trend toward diminished T-cell clonal expansion. These subtle but coordinated changes support the altered lymphocyte maturation and differentiation under conditions of CSF, which may contribute to impaired vaccine responses and protective efficacy.

## Discussion

Sleep is increasingly recognized as a critical regulator of immune function. Although clinical evidence suggests that poor sleep quality can impair vaccine–specific humoral responses, including responses to influenza vaccination (*32–37*), our understanding of the mechanisms responsible remains limited. Using a mouse model that allows for implementation of sleep fragmentation in a tightly controlled and reproducible manner (*59*), we demonstrate that CSF significantly reduces both the magnitude and quality of vaccine- induced humoral immunity, including decreased levels of influenza-specific neutralizing antibodies. These impairments are associated with transcriptional signatures indicative of altered immune cell activation and stress responses, which may compromise the coordination of T-cell–mediated support for B-cell activation and IgG class switching. These findings underscore the importance of sleep quality as a modifier of vaccine responsiveness, particularly in vulnerable populations such as shift workers, hospitalized patients and those with sleep disorders, and the elderly. While prior studies have associated poor sleep with suboptimal vaccine responses, the present study provides a high-resolution, systems-level analysis linking sleep fragmentation to molecular alternation of immune coordination.

Neutralizing antibodies are a critical correlate of protection against influenza virus infection (*60–62*), and serum antibody titers, including HAI and MN titers, are standard measures of vaccine-induced immune protection (*60, 62*). We show that CSF-exposed mice had significantly reduced HAI and MN titers compared to control mice (**Fig. 2a**). IgG binding avidity remained intact (**Fig. 2b**), suggesting that the reduced protection was due primarily to a decrease in antibody quantity rather than binding strength. CSF also reduced total IgG and class-switched subclasses (IgG1, IgG2c, and IgG3) (**Fig. 2c–d**), which are typically associated with Th1-/Th2-balanced responses (*39–41, 63*). These effects occurred despite unchanged or slightly increased number of influenza-specific antibody-secreting B-cells or IFN-γ– producing T-cells in the spleens of CSF-treated mice (**Fig. 2e**), indicating that CSF does not impair initial lymphocyte priming. Overall, these results indicate that the humoral impairments observed under CSF are not due to failure of early immune activation, but rather reflect defects in the subsequent differentiation and functional maturation of B-cells. This dissociation between early priming and final antibody output highlights the importance of sleep in regulating later stages of the adaptive immune response (*64*).

Single-cell transcriptomic profiling revealed that CSF profoundly alters B-cell developmental trajectories. Immature B-cell populations exhibited downregulation of BCR signaling, NRF2-mediated oxidative stress response, and metabolic regulatory pathways, consistent with attenuated maturation-associated signaling and reduced survival capacity during early B-cell development (**Fig. 4a**) (*65*). In contract, GC B-cells and plasma cells showed enrichment of pathways related to antigen presentation, unfolded protein response (UPR), and cellular stress, suggesting maladaptive differentiation under CSF conditions. Although class-switch recombination was clearly preserved and even elevated in GC B-cells from CSF-exposed mice (**Fig. 4b**), this was not accompanied by increased isotype expression in plasma cells or circulating antibody titers. This decoupling between CSR and effective effector output consistently suggests that CSF disrupts later stages of B-cell maturation, including plasma cell differentiation, survival, or function, thereby compromising protective humoral immunity.

Although conventional T-cell subsets (e.g., naïve, memory, Treg, and Tfh) remained transcriptionally stable by IPA analysis (**Supplementary Fig. 2c**), CSF-exposed mice showed a trend toward diminished expansion of dominant T-cell clones (**Fig. 5d**), suggesting altered repertoire architecture. A distinct DN T-cell subset exhibited selective upregulation of TCR signaling, CTLA-4 signaling, ferroptosis, and autophagy pathways, indicative of stress adaptation or chronic stimulation in this non-canonical population. While the functional consequences remain to be fully elucidated, these findings highlight CSF-induced remodeling of the T-cell compartment, particularly within subsets outside the conventional CD4⁺ and CD8⁺ lineages.

CellChat analysis revealed that CSF reconfigures B cell signaling and alters their coordination with T cells. In gSF mice, B cells upregulated ligands such as *Cd22* and MHC class I molecules, predicted to interact with *Ptprc* (CD45) and CD8 molecules on T cells and other immune subsets. These changes suggest enhanced immune activation and increased engagement with CD8⁺ T cells. However, this was accompanied by downregulation of key molecules involved in antigen presentation and costimulation, *H2-Aa* (MHC class II), *Cd86*, and *Fcer2a*, which are critical for CD4⁺ T cell–mediated B cell help, antigen-specific communication, and B cell regulation. While receptor-centered signaling networks were not significantly diminished in B cells, the overall shift indicates an imbalance between immune activation and specificity. This altered signaling landscape may reduce the quality of T–B cell interactions, compromise germinal center coordination, and contribute to the impaired antibody responses observed in CSF-exposed mice.

Together, these findings provide a systems-level view of how CSF impairs adaptive immunity by affecting the development, coordination, and repertoire programming of B- and T-cells. The implications are clinically significant. Conditions that mimic CSF, such as shift work, hospital-related sleep disruption, and sleep disorders such as obstructive sleep apnea or insomnia, are extremely common in populations that are recommended for vaccination. Alterations in sleep continuity and quality induce immune dysregulation that may significantly contribute to poor vaccine responses in these groups. Incorporating sleep assessments into vaccine protocols or optimizing vaccination timing relative to interventions aimed at sleep recovery could offer low-cost strategies to enhance immunogenicity. Moreover, our data suggest possible therapeutic avenues. Immune checkpoint modulation (e.g., PD-1/PD-L1, CTLA-4 inhibitors) may restore Tfh function and GC activity under sleep-disrupted conditions. Behavioral or pharmacologic strategies to improve sleep quality and regularity, such as cognitive behavioral therapy, melatonin supplementation, or circadian light therapy, also warrant investigation in future vaccine trials.

Future studies should investigate whether reversing CSF-induced immune changes can restore protective immunity. Longitudinal analyses of memory B-/T-cell maintenance, somatic hypermutation, and recall responses will help clarify the durability of CSF-impaired vaccine responses. The current study was conducted exclusively in young adult male mice to minimize variability from hormonal cycling and because male mice tend to be more active, which better models sleep fragmentation induced by environmental disruption; future work will include both sexes and diverse age groups to assess sex- and age-related modifiers. Ultimately, integrating sleep biology into immunization science may improve vaccine effectiveness in diverse and at-risk populations.

## Materials and Methods

### Ethical statement

The animal experiments were conducted under protocol #38742, approved by the University of Missouri’s Care and Use of Laboratory Animals in accordance with USDA Animal Welfare Regulations. All experiments were performed in a biosafety level 2 facility at the University of Missouri-Columbia under protocol #10071, in compliance with the Institutional Biosafety Committee of the University of Missouri-Columbia.

### Cells

Madin-Darby canine kidney (MDCK; CCL-34) cells were obtained from American Type Culture Collection (Manassas, VA, USA). The cells were maintained until use at 37°C under 5% CO_2_ in Dulbecco’s Modified Eagle Medium (Cat.11965084, Gibco DMEM; Thermo Fisher Scientific, Waltham, MA, USA) supplemented with 10% fetal bovine serum (Cat.10-438-026, Gibco, Thermo Fisher Scientific, Waltham, MA, USA).

### Viruses

A/California/07/2009 (HA, NA) x A/Puerto Rico/8/1934 (H1N1)pdm09 (CA/07) (BEI Resources, NR-4404) was used to prepare vaccine, and a mouse-adapted strain, A/California/04/2009 (H1N1) (CA/04-MA) (*66*), which was kindly provided by Dr. Richard Webby, was used as the challenge virus. Viruses were inoculated in 10-day-old SPF chicken embryonated eggs (VALO BioMedia North America LLC, Adel, Iowa, USA) and incubated at 37°C for 48 hours. The allantoic fluid from the infected eggs was collected for vaccine preparation or virus challenge studies.

### Vaccine preparation

The allantoic fluids were harvested and subjected to centrifugation at 3,500 × g for 30 minutes at 4°C. Polyformaldehyde inactivation was performed at a final concentration of 0.1% for 24 hours at 37°C with gentle sharking. The inactivated viruses were then centrifuge at 4 °C for 30 min at 18,000 × g in an SW28 rotor to further pellet and remove cellular debris. Pellet virions by ultracentrifugation in an SW28 rotor at 4 °C for 90 min at 112,000 × g with 5 ml chilled 30 % sucrose at bottom. The pellets were dissolved with 1 ml of phosphate buffered saline (PBS) (pH 7.4) and subjected to sucrose gradients (30%, 40%, 50%, 60%) centrifugation at 103,000 × g for 1.5 hours at 4°C. The inactivated viruses in the intermediate phase were collected and subjected to centrifugation at 103,000 × g for 1 hour at 4°C. The pellets were dissolved in sterile PBS, aliquoted and stored at -80°C until use. The hemagglutinin proteins of the vaccine were quantified by SDS-PAGE using BSA as the standard. Briefly, the vaccine was treated with PNGase F (Cat. P0704S, New England Biolabs, Ipswich, MA, USA) with the denaturing conditions for 16 hours at 37°C. The treated samples along with BSA standards were mixed with 4x Laemmli Sample Buffer and boil 10 minutes before loading the 12% SDS-PAGE. Acquired the images and analysis with ImageJ.

### Infectious virus titration

The 50% tissue culture infectious dose (TCID50) was determined using MDCK CCL-34 cells. Briefly, viral samples were serially diluted in Opti-MEM I Reduced Serum Medium (Cat. 11058021, Gibco, Thermo Fisher Scientific, Waltham, MA, USA) supplemented with 1 µg/mL TPCK-trypsin (Cat. T1426, Sigma-Aldrich, St. Louis, MO, United States). Cells were added at a density of 2 × 10^4^ cells per well in a 96-well plate with the same medium and incubated at 37°C with 5% CO_2_ for 18-20 hours. Each virus dilution (100 µL) was inoculated in MDCK cells in quadruplicate. Infected cells were fixed and then assessed for infection using ELISA assays. The number of positive and negative wells for each dilution were recorded, and the TCID50 was calculated using the method described by Reed and Muench (*67*).

### RNA extraction and qRT-PCR

Viral RNA was extracted from the tissue homogenates by using a MagMAX™ Viral/Pathogen Nucleic Acid Isolation Kit (Cat. # A48310, Thermo Fisher Scientific, Waltham, MA, USA). To quantify viral RNA copies, the viral RNAs were amplified by influenza A virus matrix gene specific qRT-PCR with the following the primers and probes: Forward 5′-GACCRATCCTGTCACCTCTGAC-3′; Reverse 5′-AGGGCATTYTGGACAAA CGTCTA-3′; and Probe 5′-[FAM]-TGCAGTCCTCGCTCA CTGGGCACG-[BHQ]-3′ (*68*). The RT-qPCR was performed by QuantStudio^TM^ 6 Flex Real-Time PCR system with Applied Biosystems^TM^ TaqMan Fast Virus 1-Step Master Mix according to the manufacturer’s protocol (Cat. 4444434, ThermoFisher Scientific, Waltham, MA, USA). The viral copy numbers in the samples were determined with the standard curve, which was generated by a plasmid containing the M gene of PR8 (*69*). The copy number determinations by qRT-PCR were performed in triplicate.

### Mouse 50% lethal dose (LD50)

LD50 for the challenge virus was determined by using 6 to 8-week-old male C57BL/6J mice from The Jackson Laboratory (Bar Harbor, ME, USA). The test virus was administered in a series of 10-fold dilution to five groups of mice (n= 5 per group), 50ul via intranasal inoculation. Mice were observed for 14 days post-administration for clinical scoring (i.e., attitude, posture, respiration, and body weight) or signs of neurologic symptoms. A mouse was euthanized when clinical score was ≥ 1.0, when neurologic symptoms (ataxia, hind-limb paralysis, incoordination) occurred, or when there was ≥ 20% weight loss. The LD50 was calculated using the Reed-Muench method, based on the mortality data observed across the different dosing groups (*67*).

### Chronic sleep fragmentation (CSF) exposures

SF was achieved by using automated SF chambers (model 80390, Lafayette instrument, Lafayette, IN, USA) which incorporate a swipe bar that moves horizontally across the cage with thins layers of bedding at set intervals. SF was performed by switching on the sweeper to a timer mode in the cage. In this mode, the sweeper requires around 7 sec to sweep the floor of the cage one way. When it reaches the very end of the cage, a relay switch engages the timer and allow 2 min before enabling the sweeper to move in the opposite direction. In between the 2 intervals, the animal is left undisturbed. On average 30 episodes of arousal from sleep per hour have been noted in patients with severe obstructive sleep apnea syndrome, which amounts to an episode every 2 min. Since our aim was to mimic as much as possible prominent sleep disruption conditions, we chose the interval of 2 min in our SF paradigm only for the light period (6am-to-6pm) corresponding to the usual predominant sleep time in rodents. The sweeper was inactive during lights-off period (6pm-to-6am). Mice had unrestricted access to food and water throughout the 24-hour cycle. Control mice resided in identical cages except that the sweeper remained immobile at all times.

### Vaccine-challenge dose optimization

We evaluated a vaccine dose per mouse ranging from 0.1 to 1.0 ug HA of A/California/07/09(H1N1) (CA/07) and the lethal challenge dose per mouse between 10 LD50 and 5000 LD50 of mouse-adapted A/California/04/09(H1N1) (CA/04-MA). Results showed that 0.1 µg HA provided 80% and 100% protection against 500 LD50 in two independent experiments, while 0.2 µg HA consistently conferred full protection (**Supplementary Fig. 1**). Doses of 0.3 µg and 1 µg HA protected against at least 5,000 LD50. Thus, in the followed study, we will use 0.2 µg HA of the CA/07 vaccine and a challenge dose of 500 LD50 of CA/04-MA.

### Evaluate influenza vaccine efficacy in CSF treated C57BL/6J mice

Six to eight-week-old male C57BL/6J mice were obtained from The Jackson Laboratory (Bar Harbor, ME USA). A total of four experimental groups were included: sleep fragmentation (SF) followed with vaccination and then virus challenge (gSF), normal-sleep control (SC) followed with vaccination and then virus challenge (gSC), PBS administration and then virus challenge (Mock), and PBS administration and then again PBS administration (Negative). SF or SC exposure lasted for 2 weeks before vaccination and continued until the end of the study (**Fig. 1a**).

During vaccination, 0.2 ug HA of inactivated vaccines or PBS in a volume of 50 µL were injected intramuscularly per mouse. Three weeks later, the mice were boosted with the same dose of vaccine or PBS. The mice were then challenged with 500 LD50 CA/04-MA or PBS in a volume of 50 µL through intranasal inoculation three weeks after the booster. Mice were observed for 14 dpi for clinical scoring (i.e., attitude, posture, respiration, and body weight) or signs of neurologic symptoms. A mouse was euthanized when clinical score was ≥ 1.0, when neurologic symptoms (ataxia, hind-limb paralysis, incoordination) occurred, or when there was ≥ 20% weight loss.

Two weeks after the booster (before the challenge), 5 mice/group were sacrificed for blood and spleen collection for immunological analyses, except for one mouse that was lost in the gSF group. Another 5 mice/group were sacrificed 3, 5, and 7 dpi to collect blood and spleens for additional immunological assessments and lung tissues for pathogenesis studies. The remaining mice/group were monitored for 14 dpi, and the same set of samples were collected. The spleen cells were obtained for transcriptomics analyses and stored in liquid nitrogen until used. Left lungs were used for pathogenicity analysis, and the right lungs for viral titration through TCID50 and qRT-PCR.

### Hemagglutination (HA), hemagglutinin inhibition (HAI) assay and Micro-neutralization (MN) assay

Sera or plasma were treated with receptor-destroying enzyme (RDE) (Cat. no. 370013, Santa Maria, CA, USA) following the manual. Briefly, sera and RDE were mixed with a ratio of 1:3 and incubated at 37°C for 20 hours before a 1-hour 56°C inactivation. PBS was then added to make a final 1:10 dilution. HA and HAI assays to test mouse sera or plasma against CA/07 were performed using 0.5% turkey erythrocytes (Cat. #7209403, LAMPIRE Biological Laboratories, Pipersville, PA) as Manual for the laboratory diagnosis and virological surveillance of influenza (*70*). MN assay to test mouse sera against CA/07 was performed with MDCK cells. The sera were 2-fold serially diluted with infection media starting at 1:10 and mixed with equal volume of 100 TCID50 viruses in 96-well plate, followed by a 1-hour incubation at 37°C. 2 × 10^4^ cells were added each well in a 96-well plate with the same medium. The plates were incubated for 18-20 hours at 37°C in 5% CO_2_ before ELISA.

### Antigen-specific B-cells by ELISPOT

100 µL of purified vaccine viruses CA/07 and HA recombinant proteins were coated on 96-well filter plates, 0.5 ug protein per well, and incubated overnight at 4°C. Plates were washed with PBS and blocked with RPMI-1640 medium (Cat. R8758-500ML, Sigma-Aldrich, St. Louis, MO, USA) containing 10% FBS (Cat. A5256801, Thermo Fisher Scientific, Waltham, MA, USA) for 1 h. 2.5 x 10^5^ cells were plated in triplicate and incubated at 37°C for 20 h. The spots were developed by ELISpot Flex: Mouse IgG (HRP) kit (Cat. 3825-2H, Mabtech, Cincinnati, OH, USA). After incubation, spleen cells will be washed and incubated with biotinylated goat anti-mouse IgG antibodies for 2 h, followed by streptavidin-conjugated HRP for 1 h. Plates were developed with TMB substrate (Cat. 3651-10, Mabtech, Cincinnati, OH, USA) for 10 min, washed, and dried. An ImmunoSpot reader (CTL, Cleveland, Ohio, USA) was used to count the developed spots in the plates. Data were expressed as the number of antibody-secreting cells (ASC) per 10^6 cells. Results were expressed as antibody secreting cells (ASC) per 10^6^ cells to quantify concentrations of IgG antibody-secreting cells.

### Virus specific T-cell ELISpot

Commercially available ELISpot Flex: Mouse IFN-γ (HRP) Mabtech ELISpot kit (Mabtech, Cat. 3321-2H, Cincinnati, OH, USA) was used to screen IFN-γ responses in mouse splenocytes at 35 dpv. We screened IFN-γ responses for CA/07 virus. The 96-well filter plates (Millipore, Cat. MSIPS4510, Sigma, St. Louis, MO, USA) were coated with 15μg/mL of purified anti-mouse IFN-γ (clone AN18; Mabtech, Cat. 3321-3-250, Cincinnati, OH, USA) in PBS at 4°C overnight. After coating, plates were blocked with RPMI1640 containing 10% FBS, L-glutamine, 1% Penicillin-Streptomycin at room temperature for 1 hour. After thawing, mouse splenocytes, 5×10^5^/well were added to coated ELISpot plates and incubated with UV inactivated vaccine strains, CA/07, prepared from a stock concentration equivalent to 25,000 TCID₅₀ per well. After incubation at 37°C, 5% CO_2_ for 72 hours plates were washed and probed with biotinylated anti-mouse IFN-γ detection mAb (clone R4-6A2; Mabtech, Cat. 3321-6-250, Cincinnati, OH, USA, 1/1000 diluted in PBS with 0.5% FBS) followed by streptavidin-conjugated HRP (Mabtech, Cat. 3310-9, Cincinnati, OH, USA, 1/1000 diluted in PBS with 0.5% FBS and TMB substrate (Mabtech, Cat. 3651-10, Cincinnati, OH, USA). After air drying, distinct spots were visualized and scanned using an ImmunoSpot reader (CTL, Cleveland, Ohio, USA). Background (without stimulus) was subtracted, and data were expressed as the number of spot-forming units (SFU) per 10^6^ cells.

### Immunohistochemistry staining and pathological analysis of lungs after viral infection

The left lungs from mice after euthanasia were fixed in 10% neutral buffered formalin solution. To evaluate the quantity of influenza antigen present in the mouse lung, slides were incubated with NP rabbit monoclonal antibody (Cat. # MA5-42363, Invitrogen, Waltham, MA, USA) overnight at 4°C after blocking with normal horse serum. Sections were treated with VR Horse Anti-Rabbit IgG Polymer Detection Kit (Cat. #MP-6401-15, Vector Laboratories, Newark, CA, USA) according to the manufacturer’s instructions. DAB (Cat. #SK-4105, Vector Laboratories, Newark, CA, USA) was incubated with slides for 5 minutes for color development. Sections were counterstained with hematoxylin, washed, dehydrated, and covered with cover slips. The stained slides were scanned at a 40x magnification by MU Pathology Informatics Whole-Slide Imaging Analytics Lab with Aperio ScanScope CS. Influenza NP-positive cells were quantified using positive cell detection function from QuPath 0.4 (*71*) across the entire slide, the H-score is evaluated and compared among groups (*72*).

### Antigen-specific total IgG and subclass IgG detection using ELISA

Nunc MaxiSorp microtiter plates (Thermo Fisher Scientific, Cat. 439454, Waltham, MA, USA) were coated with 0.5 µg of purified CA/07 in phosphate-buffered saline (PBS) at 4°C overnight. Plates were then blocked at room temperature for 1 hour with blocking buffer (PBS containing 0.1% Tween 20 and 5% Blotting-Grade Blocker, Cat. 1706404, Bio-Rad, Hercules, California, USA). Mouse plasma was 2-fold serially diluted in blocking buffer starting at 1:100 and incubated at room temperature for 1 hour. For different antibody subclasses detection, secondary antibodies of each conjugated with HRP were applied and incubated at room temperature for 1 hour. Goat anti-Mouse IgM Secondary Antibody (Cat. 31440, Invitrogen, Waltham, MA, USA), Goat anti-Mouse IgG Fc Secondary Antibody (Cat. 31437, Invitrogen, Waltham, MA, USA), Goat Anti-Mouse IgG3 (Cat. 75952S, Cell Signaling Technology, Danvers, MA, USA), Goat Anti-Mouse IgG2b (Cat. 43593S, Cell Signaling Technology, Danvers, MA, USA), Goat Anti-Mouse IgG1 (Cat. 96714S, Cell Signaling Technology, Danvers, MA, USA), Goat Anti-Mouse IgG2c (Cat. 56970S, Cell Signaling Technology, Danvers, MA, USA). Color development was achieved by adding Pierce™ 1-Step™ Ultra TMB-ELISA Substrate Solution (Thermo Fisher Scientific, Cat. 34029, Waltham, MA, USA) for 10 minutes, and the reaction was stopped by adding 50 µL of 1N sulfuric acid (Cat. SA8184, Fisher Chemical, Thermo Fisher Scientific, Waltham, MA, USA) diluted in Milli-Q water. Optical density at 450 nm (OD450) was measured using a BioTek Cytation 5 Cell Imaging Multimode Reader (Agilent, Santa Clara, CA, USA). The area under the entire antibody titration curve was calculated. Briefly, the plots of OD450 values vs log of the sera dilution in the ELISA experiment were used to obtain the values of area under curve (AUC) by GraphPad Prism (version 10.1.2).

### Antigen-specific antibody avidities

For avidity assays, after the sera incubation, the plates were washed and overlaid with 100 µL/well of 6 M urea for 15 minutes, followed by thorough washing. Bound antibodies were detected using peroxidase-conjugated goat anti-rabbit IgG (Cat. A16110, ThermoFisher, Waltham, MA, USA). Color development was achieved by adding Pierce™ 1-Step™ Ultra TMB-ELISA Substrate Solution (Thermo Fisher Scientific, Cat. 34029, Waltham, MA, USA) for 10 minutes, and the reaction was stopped by adding 50 µL of 1N sulfuric acid (Cat. SA8184, Fisher Chemical, Thermo Fisher Scientific, Waltham, MA, USA) diluted in Milli-Q water. Optical density at 450 nm (OD 450) was measured using a BioTek Cytation 5 Cell Imaging Multimode Reader (Agilent, Santa Clara, CA, USA). The avidity index was calculated based on the area under the entire antibody titration curve (*73*). Briefly, the plots of OD values vs log of the sera dilution in the ELISA experiment with and without urea treatment were used to obtain the values of area under curve (AUC) by GraphPad Prism (version 10.1.2). The antibody index equals to an ACU ratio of treated and untreated reactions. All samples were assessed in duplicate and mean titers will be calculated (*74*).

### Single-cell RNA and TCR/BCR sequencing

To analyze B- and T-cell transcriptional responses to influenza vaccination, we performed paired scRNA-seq and scTCR/BCRseq on spleen cells from mice in gSF (n=3) and gSC (n=3). Single-cell cDNA libraries with TCR ab and BCR light heavy chains paired VDJ libraries were prepared using the 10x Chromium Next GEM Single Cell 5′ reagent kit (v2) according to the manufacturer’s instructions. Sequencing was performed on the S4 flow cell of the NovaSeq 6000 sequencer (Illumina) to obtain paired end reads. Sequence reads were aligned to the mouse reference dataset GRCm39, followed by creation of barcode gene matrices using Cell Ranger v8.0.0 (10x Genomics). Clustering analyses were performed using Seurat (v4.4.0) (*75*). To filter out low-quality genes and cells, only genes expressed in more than 3 cells and excluding cells with aberrantly high or low gene counts and high mitochondrial gene expression. Scrublet (v.0.2.3) was used to detect doublets (*76*). Afterwards, transcript counts were normalized and log2 transformed. Highly variable genes were used to produce the principal component analysis (PCA). Dimensionality reduction uniform manifold approximation and projection (UMAP) and Louvain clustering were carried out. Cell lineages were manually annotated based on known markers genes found in the literature, in combination with algorithmically defined DEGs in each cluster, using *FindAllMarkers* function with the Wilcoxon test. Then, the partition-based graph abstraction (PAGA) for trajectory analysis was performed using the *scanpy.tl.paga* function from Scanpy (v1.9.1) (*77*), which reconciles clustering and pseudotemporal ordering algorithms and allows the inference of complex cell trajectories and differentiation trees (*78*). The DEGs between treatments in each cluster were computed using *FindMarkers* with the Wilcoxon test (logfc.threshold = 0.25, min.pct = 0.1). The R package EnhancedVolcano (v1.14.0) (https://github.com/kevinblighe/EnhancedVolcano) was used to visualize the DEGs in the selected clusters. Cell-cell interaction from scRNA-seq data was predicted using R package CellChat (v2.1.1) based on the expression of known ligand-receptor pairs in mice (*54*). A comparison analysis was performed between treatments. The Interaction circular maps of upregulated or downregulated interactions were built using the function *netVisual_chord_gene*. The size of the interaction arrow is in accordance with the transcriptional level of ligand or receptor genes in each cell type.

Single-cell TCR and BCR sequences were aligned and quantified using cellranger vdj (10x Genomics) pipeline against the GRCm38 mouse VDJ reference genome. Filtered annotated contigs for TCRs and BCRs were analyzed using the Scirpy package (v0.10.1) (*79*), after which TCR and BCR data were integrated with gene expression data. Cells without a full TCR ab chain or BCR light and heavy pairs were excluded.

Clonotypes were defined based on CDR3 nucleic acid sequence identity and clonal size that were estimated using scirpy.tl.define_clonotypes and scirpy.tl.clonal_expansion, respectively. Clone diversity curves that measured Hill’s diversity metric across diversity orders (q) from 0 to 4 were created using the R package alakazam (v1.2.1) with the alphaDiversity function (*80*).

### Ingenuity Pathway Analysis (IPA)

Differentially expressed genes (DEGs) were identified based on an adjusted p-value < 0.05 and a log2 fold change > 1. The list of DEGs, including Ensembl gene IDs, avg_log2FC, fold changes, p-values, and adjusted p-values, was uploaded to the Ingenuity Pathway Analysis (IPA) platform (QIAGEN Inc., https://digitalinsights.qiagen.com/products-overview/discovery-insights-portfolio/analysis-and-visualization/qiagen-ipa/). Core Analysis function was used to identify canonical pathways significantly associated with the DEGs, applying a p-value threshold of 0.05. The analysis was configured to focus on mouse, using the Ingenuity Knowledge Base as the reference set. Confidence levels were set to "experimentally observed," and all available data sources were included.

Following core expression analyses, upstream regulator analysis was performed to predict transcription factors and signaling molecules driving the observed gene expression changes, with z-scores > 2 or < -2 indicating activation or inhibition. Pathways and genes were considered significant if they had a -log(p-value) greater than 1.3 (p-value < 0.05) and a z-score greater than 2 (indicating pathway activation) or less than -2 (indicating pathway inhibition). Significantly enriched canonical pathways, particularly those related to the immune system, were identified, and visualized through IPA to assess their biological relevance to the study.

## Data availability

We have submitted the scRNAseq data collected from this study to GenBank under the BioProject accession number PRJNA XXX.

## Statistical analysis

All statistical analysis was performed by GraphPad Prism 9.0.1 software (GraphPad software, La Jolla, Ca). The nonparametric unpaired Mann–Whitney U test was used to compare serological titers, including HAI, MN, and antibody binding avidity, across experimental groups. The nonparametric unpaired Kruskal–Wallis test was used to compare H-scores among groups. Unpaired Student’s t-tests were applied to assess differences in the CDR3 length of B-cells and expanded clones in T-cells. P-values <0.05 were considered significant.

## Acknowledgments

The authors thank Stephanie Nava, George Sarafianos, Rebecca Patterson, Mingyi Zhou, and Darling Melany De Carvalho Madrid for their assistance in animal study and Kritika Prasai for her assistance in data visualization. We would like to acknowledge Samantha Gerb and Sarah N. Schlink for their veterinary support. We also thank the University of Missouri Genomics Technology Core for assistance with scRNA-seq data acquisition and the Proteomics Core for support with mass spectrometry analyses. This grant was supported by the startup fund from University of Missouri.

---

**Supplementary Fig. 1.**
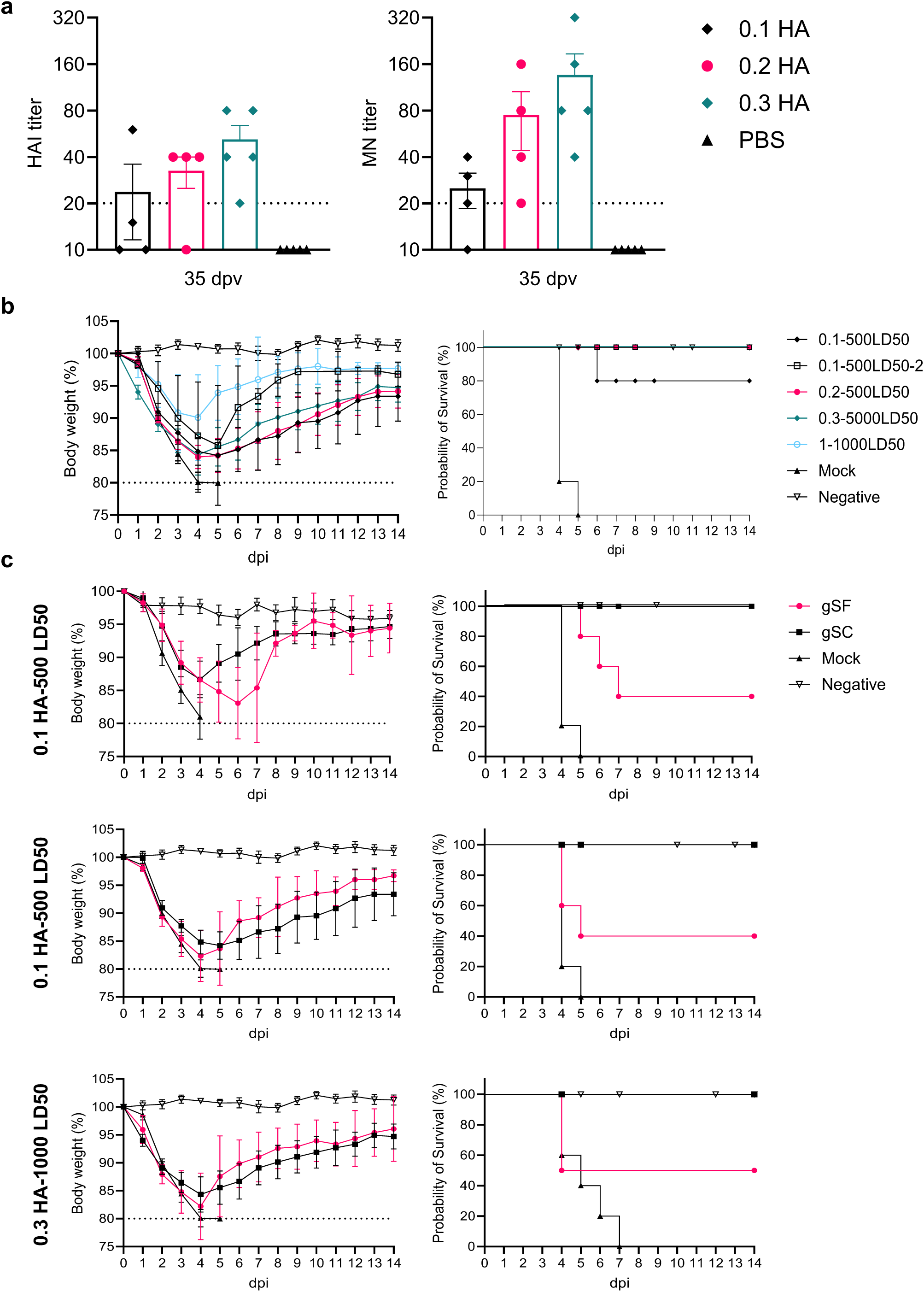
Dose optimization for influenza vaccination and viral challenge in mice. **a)** Hemagglutination inhibition (HAI) and microneutralization (MN) titers in mice immunized intramuscularly with 0.1, 0.2, or 0.3 µg of inactivated A/California/07/2009 (H1N1) (CA/07) hemagglutinin (HA) antigen, or PBS as control. **(b)** Body weight loss and survival rates for mice vaccinated with varying doses (0.1, 0.2, 0.3, and 1.0 µg) of inactivated CA/07 HA antigen or PBS, followed by lethal challenge with 500–5000 LD_50_ of mouse-adapted CA/04 (CA/04-MA) virus. **c)** Body weight loss and survival rates from three independent experiments using different vaccine–challenge dose combinations: 0.1 µg HA – 500 LD_50_, 0.1 µg HA – 500 LD_50_ (repeat), and 0.3 µg HA – 5,000 LD_50_. Experiment 1: 0.1 µg HA – 500 LD_50_; gSF (n = 9), gSC (n = 10), Mock (n = 7), Negative (n = 9); Experiment 2: 0.1 µg HA – 500 LD_50_; gSF (n = 5), gSC (n = 5), Mock (n = 5), Negative (n = 5); Experiment 3: 0.3 µg HA – 5000 LD_50_; gSF (n = 5), gSC (n = 5), Mock (n = 5), Negative (n = 4). Mice were housed under standard conditions in ABSL-2 containment. CSF was induced in gSF groups beginning 14 days prior to vaccination and continued through challenge, while gSC groups maintained normal sleep. Mock groups received PBS in place of vaccine, and negative controls were neither vaccinated nor challenged.

**Supplementary Fig. 2.**
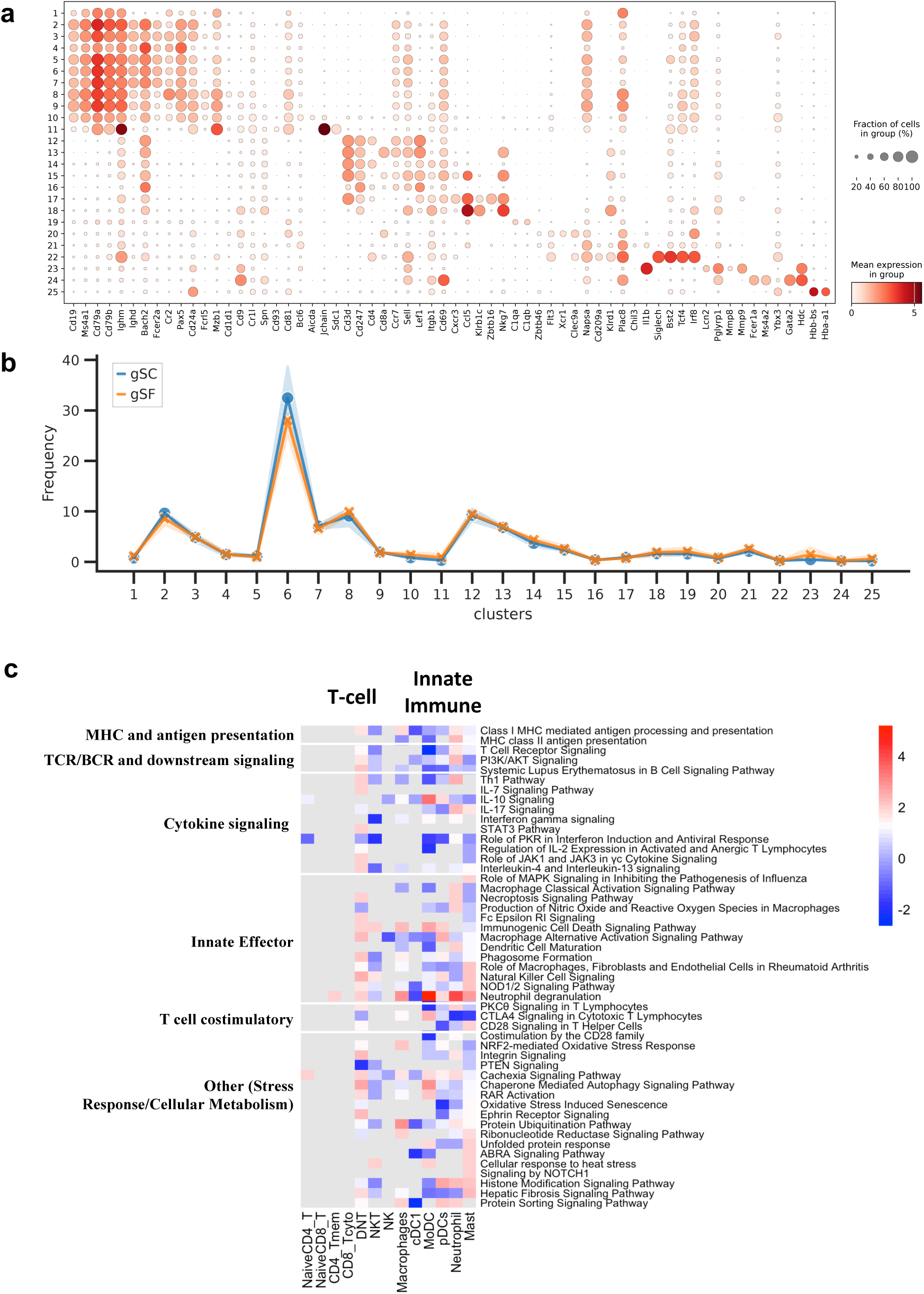
Single-cell analyses of innate immune cells and T-cells following CSF exposure. **a)** Canonical marker gene expression across splenic immune cell clusters. Dot plot showing the expression of canonical lineage and functional marker genes across 25 transcriptionally defined cell clusters identified by single-cell RNA sequencing of splenocytes from normal-sleep control (gSC) and CSF-exposed (gSF) mice. Each row corresponds to a distinct cell cluster, and each column represents a selected marker gene. Dot size reflects the percentage of cells in each cluster expressing the gene, while color intensity indicates the mean expression level of the gene within that cluster (see scale bar). Marker genes were used to assign cluster identities to major immune subsets, including B-cells (e.g., Cd19, Ms4a1, Cd79a, Ighm), T-cells (e.g., Cd3e, Cd4, Cd8a), dendritic cells (e.g., Itgax, Cd209a), macrophages and monocytes (e.g., Cd68, Lyz2, Adgre1), NK cells (e.g., Nkg7, Klrb1c), neutrophils (e.g., S100a8, S100a9), and plasma cells (e.g., Synd1, Jchain, Prdm1). Among six FoB cells, Cluster 3 is GC B-cell precursors (Ighd-, Cd38-, Fas+) , Cluster 4 is early activated FoB cells (Ccr7+, Cxcr5+, Cd86+, Cd83+, Bcl6+, Cr2+, Irf4+, Mki67+), and Cluster 5, 6, 7 are resting FoB cells (or recirculating FoB cells)(high IgD, low IgM, Thy1+). **b)** Distribution of immune cell cluster frequencies in CSF and normal-sleep control mice. Line plot showing the relative frequencies of 25 transcriptionally defined immune cell clusters in splenocytes from CSF-exposed (gSF, orange) and control (gSC, blue) mice. Frequencies are expressed as the percentage of total cells assigned to each cluster. Data points and lines indicate group means, and shaded areas denote standard deviation. Overall, cluster distributions were broadly conserved between groups, indicating comparable cellular composition despite transcriptional reprogramming in select subsets. **c)** Ingenuity Pathway Analysis (IPA) of T-cell and innate immune subsets. Analyses were performed using the QIAGEN IPA platform. Core Analysis function was used to identify canonical pathways significantly associated with the DEGs, applying a p-value threshold of 0.05. Pathways were considered significant if they met a –log(p-value) > 1.3 (equivalent to p < 0.05) and a Z-score > 2 (activation) or < –2 (inhibition). Only pathways with a Z-score ≥ 2 or ≤ –2 in at least one cell type were included (performed using R packages). Positive Z-scores (red) indicate pathway upregulation in CSF-treated mice. Pathways were grouped into six categories: 1) MHC and antigen presentation, 2) TCR/BCR and downstream signaling, 3) Cytokine signaling, 4) Innate effector functions, 5) T-cell costimulatory signals, 6) Other such as stress response and cellular metabolism.

**Supplementary Fig. 3.**
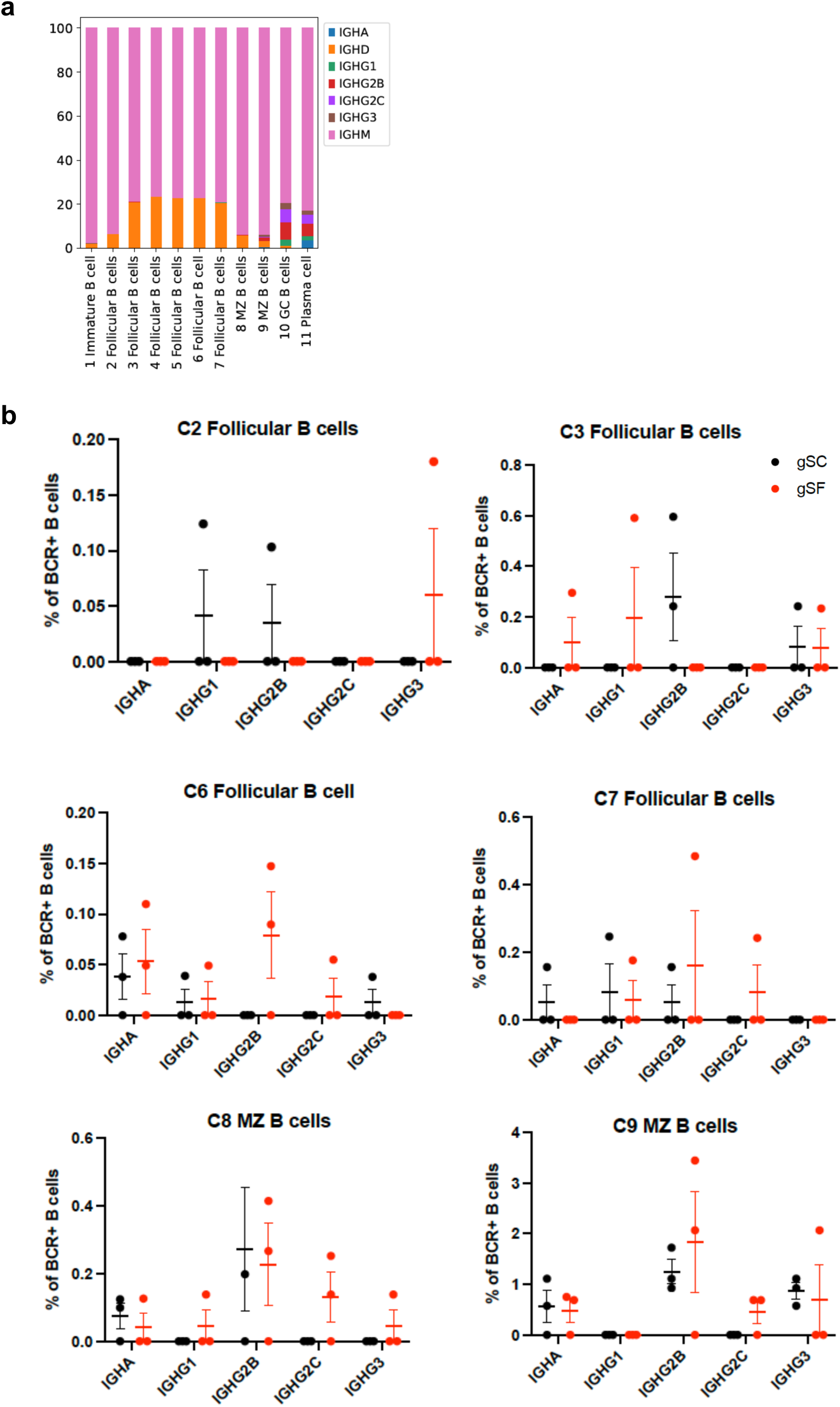
Distribution of immunoglobulin isotypes across B-cell subsets in CSF and normal-sleep control mice. **a)** Stacked bar plot showing the relative abundance of immunoglobulin isotypes (IGHM, IGHD, IGHG1, IGHG2B, IGHG2C, IGHG3, and IGHA) among BCR⁺ cells across transcriptionally defined B-cell subsets. Immature B-cell cluster (C1), Follicular B-cell clusters (C2–C7), marginal zone (MZ) B-cells (C8–C9), germinal center (GC) B-cells (C10), and plasma cells (C11) are shown. Most early B-cell subsets (e.g., immature and follicular B-cells) are dominated by IGHM, whereas GC and plasma cells show greater isotype diversity. **b)** Dot plots show the percentage of BCR⁺ cells expressing each isotype (y-axis) in individual follicular B-cell clusters and marginal zone B-cells, comparing CSF-exposed (gSF, red) and normal-sleep control (gSC, black) mice. Each dot represents one mouse; bars indicate mean ± SEM.

